# Motor Neurons Decode Cholinergic Inputs via Spatially Distinct nAChR Subunits to Drive Locomotion in Drosophila larvae

**DOI:** 10.1101/2025.06.07.658461

**Authors:** Ankura Sitaula, Lizzy Olsen, Arianna Mogharrabi, Aref Zarin

## Abstract

Neural circuits consist of neurons that differ not only in their neurotransmitter identities but also in the types and subcellular localization of neurotransmitter receptors (NRs) they express. This receptor diversity enables distinct responses to the same neurotransmitter, highlighting the need to understand NR distribution and function to fully interpret circuit logic. Here, we focus on nicotinic acetylcholine receptors (nAChRs), the primary mediators of fast excitatory transmission in the *Drosophila* central nervous system (CNS). Functional nAChRs are pentamers assembled from a pool of 10 subunits (α1–α7, β1–β3), yet their in vivo expression and function remain poorly defined.

We used T2A-Gal4 lines and endogenous protein tagging to examine nAChR expression in larval motor neurons (MNs) and identified eight subunits (α1–α3, α5–α7, β1, β2) expressed in these cells. MN-specific knockdown of individual subunits caused distinct locomotor defects, indicating their functional importance. Co-localization analysis revealed some subunit pairs are frequently co-expressed at the same synapses, while others localize to distinct subcellular domains. Supporting this, double knockdown of co-localized subunits did not worsen locomotor phenotypes compared to single knockdowns, whereas knockdown of non-co-localized subunit pairs produced additive defects.

These results suggest that different nAChR subtypes are strategically positioned in discrete synaptic domains within single MNs, where they serve non-redundant roles. Our findings provide new insight into the spatial organization and functional diversity of nAChRs in motor circuits that drive locomotion.

**Significance Statement:** Motor circuits rely on precise neurotransmitter signaling, yet the diversity and subcellular organization of neurotransmitter receptors remain poorly understood. Using Drosophila larvae, we show that motor neurons express multiple nicotinic acetylcholine receptor (nAChR) subunits, which localize to distinct synaptic domains and play non-redundant roles in locomotion. These findings reveal a previously underappreciated level of receptor compartmentalization within single neurons and demonstrate that spatially organized nAChRs are essential for coordinated movement. By integrating genetic, imaging, and behavioral approaches, our work provides a new framework for understanding how receptor diversity shapes motor output and highlights the importance of mapping receptor localization to decode circuit function in both health and disease.

## Introduction

With advances in electron microscopy-based connectomics, we now have detailed wiring diagrams of neural circuits across species (1-4). These circuits comprise neurons with diverse neurotransmitter identities, each potentially expressing a broad array of neurotransmitter receptors (NRs) with distinct properties and subcellular localizations. This receptor diversity enables circuits to generate varied responses to the same neurotransmitter. To fully interpret connectomes and understand circuit function, it is essential to map the distribution and roles of NRs within these networks. To address this gap, we leverage the powerful genetic tools and high-resolution EM connectome available for *Drosophila* motor circuits to investigate the diversity and function of nAChRs in larval MNs.

Acetylcholine (ACh) acts through both ionotropic and metabotropic receptors to mediate fast and slow signaling, respectively. nAChRs, the ionotropic ACh receptors, are members of the cys-loop superfamily of pentameric ligand-gated ion channels and serve as central components of the cholinergic system (5). A defining feature of nAChRs is their molecular complexity, which gives rise to extensive structural and functional diversity. In *Drosophila*, 10 nAChR subunits—seven α and three β—are encoded, compared to 10 α and four β subunits in vertebrates (6-12). These subunits assemble into homo- or heteropentameric cation channels that mediate fast excitatory transmission. Each mature pentamer likely exhibits distinct expression patterns, channel properties, and regulatory mechanisms. However, the functional logic of this subunit diversity—and how it contributes to receptor heterogeneity within neural circuits—remains poorly understood.

Disruptions in the cholinergic system have been implicated in a range of human neurological and psychiatric disorders, including Alzheimer’s and Parkinson’s diseases, depression, autism, ADHD, epilepsy, and schizophrenia (13-15). As central components of this system, nAChRs are promising targets for drug development aimed at treating these conditions (15). In addition, insect nAChRs are the primary targets of widely used agricultural insecticides (8, 10, 12), though resistance to these compounds has emerged in many insect populations (16). A deeper understanding of nAChR expression and function is therefore critical not only for advancing therapeutic strategies for human diseases but also for designing more selective and sustainable insecticides with minimal off-target effects on beneficial species, including humans. While heterologous systems such as *Xenopus laevis* oocytes have been invaluable for studying nAChR structure and function, they do not fully capture receptor behavior in native neuronal environments (17-19). Receptors expressed in these systems may differ from endogenous receptors due to differences in lipid composition, pre- and post-translational modifications, and codon usage, all of which can influence receptor localization, assembly, and function (17). As a result, direct in vivo studies remain essential yet scarce, leaving key aspects of nAChR biology unresolved.

*Drosophila* offers several key advantages for studying the distribution and function of nAChRs in the nervous system. Powerful genetic tools now allow endogenous tagging of nAChR subunits within their native genomic context (10, 20-23), avoiding artifacts associated with overexpression systems, such as aberrant protein accumulation or mislocalization. In parallel, high-resolution EM connectomes for both larval and adult *Drosophila* provide an ideal framework for mapping endogenously tagged nAChR subunits onto known neural circuits (24-27). nAChR facilitate excitatory neurotransmission within the *Drosophila* brain and are essential for neural development, synaptic plasticity, and cognitive activities such as learning and memory (28). Like their mammalian counterparts, *Drosophila* nAChRs also regulate dopamine release, supporting a conserved role in dopaminergic signaling (29, 30). Importantly, individual subunits contribute to specific biological processes. Temporal regulation of α1, α5, α6, and α7 supports maturation of cholinergic synapses (23, 31, 32). Null mutants of several nAChR subunits display diverse behavioral and morphological phenotypes, indicating pleiotropic roles beyond neurotransmission. For example, α1, α2, α3, and α6 null mutants exhibit reduced climbing ability and decreased adult longevity, while α1, α2, α5, and β3 mutants show curled abdomens, and α1, β1, and β3 mutants exhibit defective wing inflation (10, 33-35). Additionally, α7 is essential for the escape reflex via the giant fiber system (36), and α1 and α3 are involved in complex behaviors such as courtship and sleep (37-39). As these null mutants lack receptor expression throughout the animal, tissue-specific manipulations are critical to uncover subunit functions within defined neural populations. Together, these findings highlight the diverse roles of nAChRs in behavior, yet their circuit-level functions—particularly within motor circuits driving locomotion—remain largely unexplored.

To address these questions, we performed genetic studies in *Drosophila* larval MNs, which receive excitatory inputs from several cholinergic pre-motor neurons (PMNs) (24, 40-43). Using a suite of genetic tools—including T2A-Gal4 lines, endogenous tagging, and conditional tagging of nAChR subunits—we identified a diverse array of nAChR subunits expressed in larval MNs and mapped their distribution at single-neuron resolution. MN-specific RNAi knockdown revealed that many subunits are essential for normal locomotion, as their depletion caused significant crawling defects affecting peristaltic efficiency, duration, protopodium folding, and displacement. Co-localization analysis of endogenously tagged nAChR subunit pairs, combined with single and dual knockdown experiments, revealed distinct patterns of subunit distribution within motor dendrites. While some subunits co-localized at the same synapses, others were spatially segregated, indicating the existence of synapse-specific receptor profiles. Together, these findings underscore the essential role of nAChR-mediated excitation in MN function and demonstrate that MNs depend on a diverse and spatially organized repertoire of nAChRs, with minimal functional redundancy among subunits.

## Results

### nAChR Subunits Are Widely Expressed in Larval Motor Neurons

The Drosophila larval body is composed of three thoracic segments (T1–T3) and nine abdominal segments (A1–A9), with the corresponding ventral nerve cord (VNC) organized into 12 segmentally aligned neuromeres (**Fig. 1A**). The larval CNS consists of two brain lobes and a segmented VNC, which functions analogously to the vertebrate spinal cord. Each VNC segment controls the muscles of one body segment (**Fig. 1B**). Each abdominal segment contains approximately 30 bilaterally symmetrical muscle pairs (**Fig. 1C**), innervated by a similar number of glutamatergic excitatory motor neurons (MNs) projecting from the corresponding VNC segment (44-46). Larval MNs are broadly categorized into two classes: phasic Type Is MNs, which typically innervate multiple muscles and form small synaptic boutons, and tonic Type Ib MNs, which target one to three muscles and form larger boutons at the neuromuscular junction (NMJ) (47-59).

**Figure 1.**
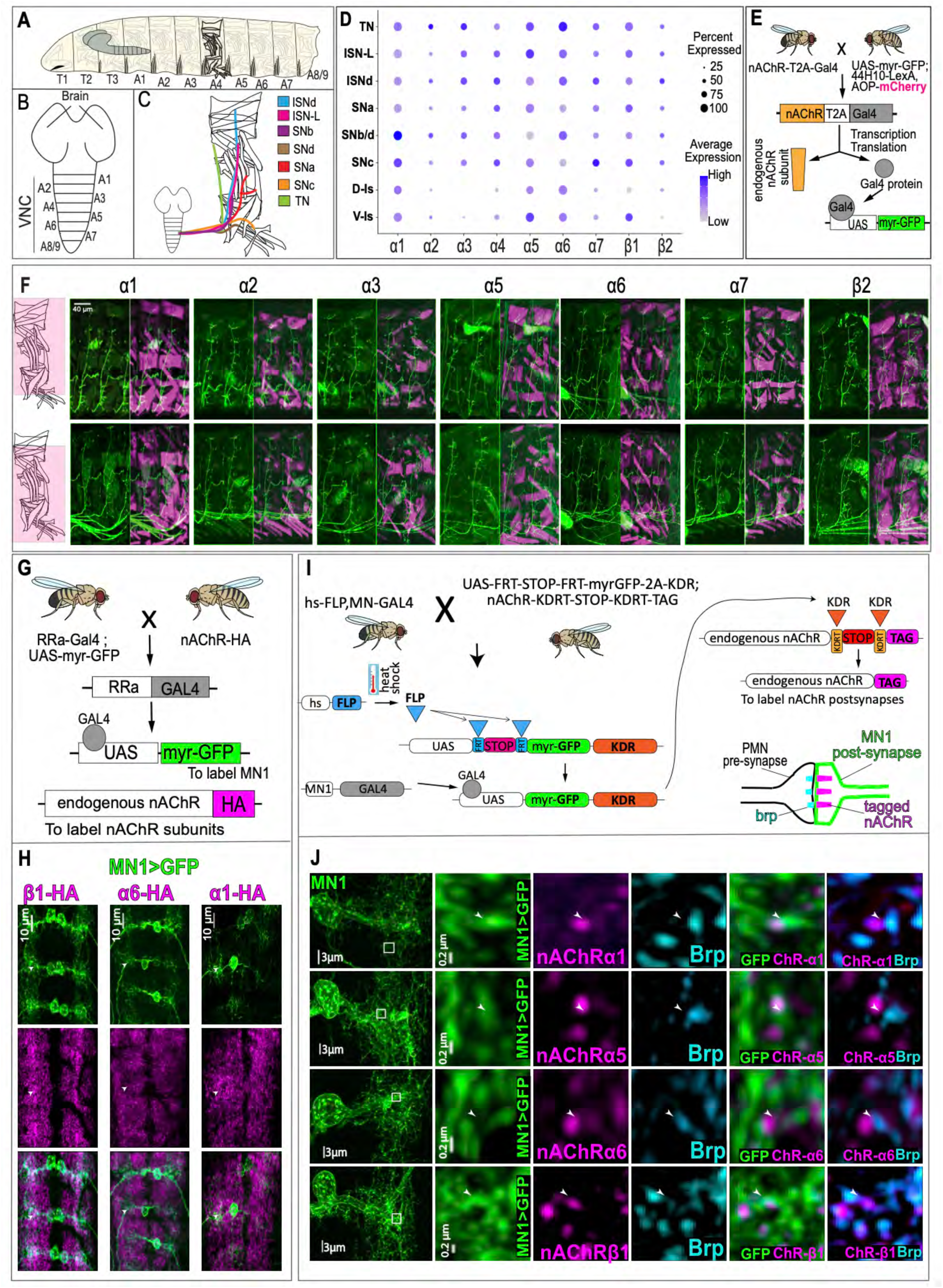
Larval MNs Co-express a Diverse Repertoire of nAChR Subunits. (A) Schematic of the Drosophila larval body, composed of three thoracic and nine abdominal segments. (B) Anatomy of the CNS, showing two brain lobes and the segmented ventral nerve cord (VNC) controlling body segments. (C) Each abdominal (A1–A6) half-segment contains ∼30 muscles, innervated by ∼30 excitatory glutamatergic motor neurons (MNs) projecting via distinct nerve routes (color-coded): ISNd, ISNL, SNa, SNb, SNc, SNd, and TN. (D) Dot plot showing nAChR subunit expression across MN clusters, based on scRNA-seq data from Nguyen et al. (62). Dot size reflects the percentage of expressing cells; color intensity reflects average expression level. (E) nAChR-T2A-Gal4 reporter system schematic: T2A-mediated ribosomal skipping enables independent translation of Gal4 and endogenous nAChR subunit. UAS-myr-GFP visualizes nAChR-expressing cells; muscles are labeled with GMR44H10-driven mCherry. (F) Expression patterns of seven nAChR subunits (α1, α2, α3, α5, α6, α7, β2) in MNs, visualized using the strategy in (E). (G) Cross strategy to express myr-GFP in MN1 and endogenously tag nAChRs. (H) HA-tagged α1, α6, and β1 subunits colocalize with MN1 dendrites, visualized using the strategy in (G). (I) Conditional tagging strategy using FLP- and KDR-mediated recombination to selectively tag nAChRs in MN1; Bruchpilot (Brp) marks presynaptic sites. (J) Endogenous conditional tagging reveals clustered nAChR puncta adjacent to Brp-positive active zones in MN1 dendrites, visualized using the strategy in (I). Scale bars as indicated. Genotypes are provided in the SI Appendix.

All larval MNs receive excitatory cholinergic input from premotor neurons (pre-MNs), indicating that cholinergic receptors are essential components of the MN postsynaptic compartment (24, 60, 61). Recent single-cell RNA sequencing (scRNA-seq) of the larval VNC further supports this (62), showing widespread expression of nAChR subunit transcripts across MNs (**Fig. 1D**). However, expression of individual subunits at the level of single, identified MNs has not been systematically characterized.

To address this gap, we utilized T2A-Gal4 lines for each nAChR subunit (63-65) in combination with UAS-membrane-targeted myristoylated GFP (myr-GFP) to visualize subunit-expressing MNs. Postsynaptic muscles were simultaneously labeled using 44H10-LexA > LexAop-mCherry, enabling identification of individual MNs based on their target muscle(s) (**Fig. 1E**). This strategy allowed tracing of motor axon terminals and NMJs in intact larvae, enabling determination of subunit expression at single-MN resolution. Using this approach, we found that at least seven nAChR subunits (α1–α3, α5–α7, and β2) are expressed in the majority of larval MNs (**Fig. 1F**). For example, MN1, which innervates muscle 1, expresses all seven of these subunits based on nAChR-T2A-Gal4 > myr-GFP labeling. These findings demonstrate broad and combinatorial expression of nAChR subunits across identified MNs, suggesting that multiple receptor subtypes may contribute to cholinergic signaling within the motor system.

To corroborate these findings, we examined nAChR subunit expression in MN1 using endogenously HA-tagged lines for α1, α6, and β1 generated in previous studies (20, 23). These lines were crossed to RRa-Gal4 > UAS-myrGFP to label MN1 (**Fig. 1G**). Numerous HA-positive puncta were observed within the MN1 dendritic arbor for all three subunits, confirming their expression at the protein level (**Fig. 1H**).

To further refine subunit localization in a cell-type-specific manner, we employed a conditional tagging strategy developed by Sanfilippo et al. (20), in which nAChR subunits are epitope-tagged only when their endogenous alleles are transcriptionally active in the cell of interest (**Fig. 1I**). This tool, currently available for α1, α5, α6, and β1, revealed OLLAS- or HA-tagged puncta in MN1 dendrites for all four subunits. Notably, these puncta were frequently juxtaposed to presynaptic active zones labeled by Bruchpilot (Brp), suggesting that these nAChR subunits are incorporated into postsynaptic specializations of MN1 (**Fig. 1J**).

Together, data from T2A-Gal4 expression analysis, endogenous HA tagging, conditional OLLAS tagging, and scRNA-seq collectively confirm that multiple nAChR subunits are expressed in larval MNs and are synaptically localized. These complementary approaches provide high spatial resolution for identifying subunit expression at the level of single, identified MNs. Given their widespread and synaptic distribution, we hypothesize that these nAChR subunits play critical roles in mediating cholinergic input and are essential for proper motor circuit function.

### MN-Specific Knockdown of Individual nAChR Subunits Impairs Peristalsis Efficiency, Duration, and Protopodium Dynamics During Crawling

Given the widespread expression of nAChR subunits in larval MNs, we hypothesized that these receptors mediate excitatory cholinergic input and are essential for motor circuit function. To test this, we used the pan-MN driver OK6-Gal4 to perform UAS-RNAi–mediated knockdown of each of the ten Drosophila nAChR subunits. We focused on forward crawling, the primary locomotor behavior in larvae used for exploratory foraging. This behavior relies on the coordinated activation and inactivation of segmentally paired muscles, propagating peristaltic contractions from posterior to anterior segments.

As a behavioral readout, we quantified forward crawling by measuring the length traveled over 60 seconds (**Fig. 2A**). Knockdown of eight out of ten subunits (α1–α6, β2, and β3) significantly reduced crawling length, indicating their critical roles in MN function (**Fig. 2B** and **Movie S1**). To ensure these phenotypes were not due to off-target RNAi effects, we repeated knockdown experiments for α3, α5, and α6 using independent non-overlapping RNAi lines, which reproduced the observed defects (**SI Appendix, Fig. S1**). Although α7 and β1 were expressed in MNs, their knockdown had no significant effect on crawling behavior, even when tested with multiple RNAi lines (**SI Appendix, Fig. S1**). This suggests that these subunits may be functionally redundant or dispensable in this context.

**Figure 2.**
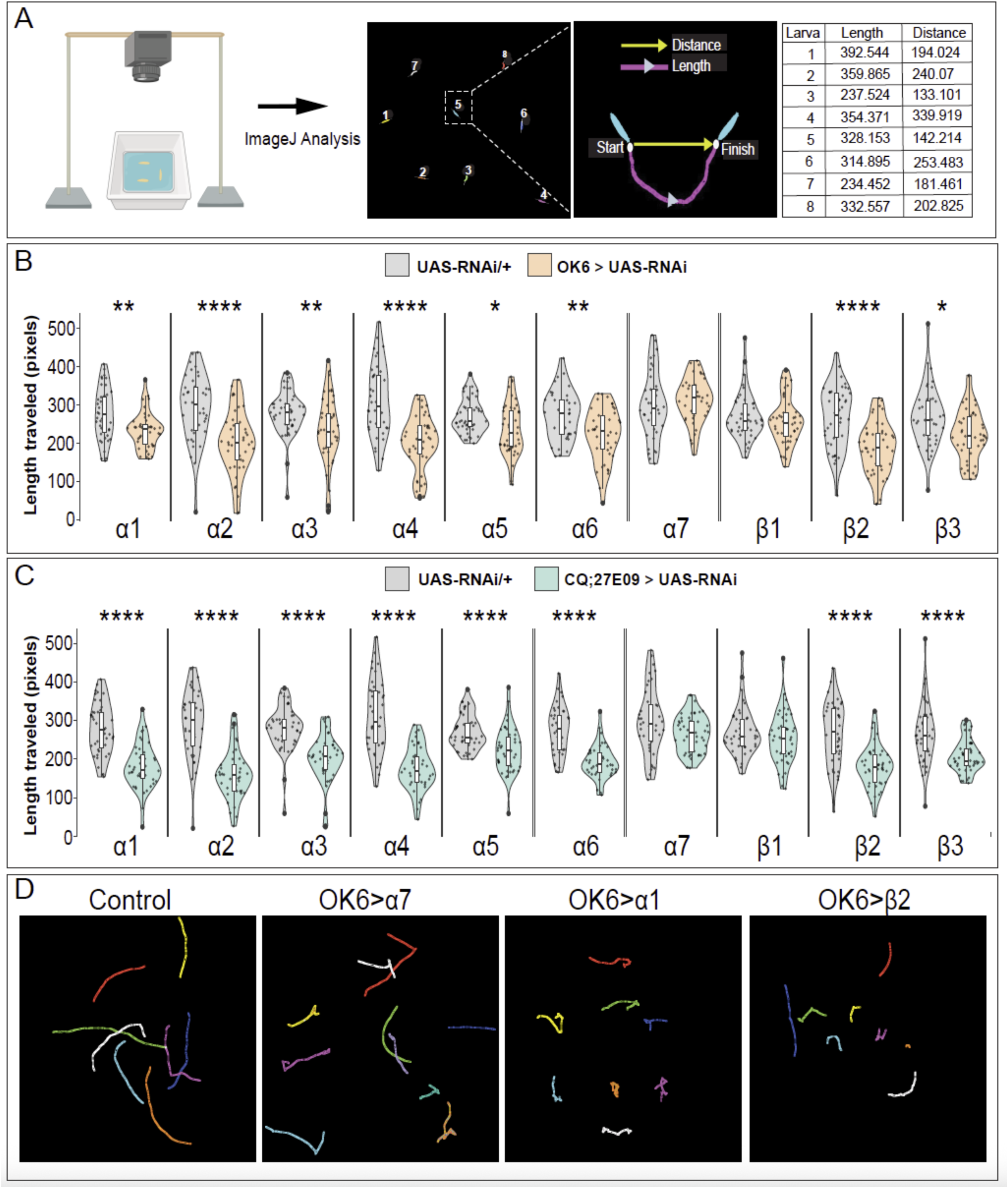
Single nAChR Subunit Knockdown in Motor Neurons Impairs Larval Locomotion. (A) Workflow for ImageJ and wrMTrck analysis of larval locomotion. Larvae were numbered and tracked to measure path length. (B) MN-specific knockdown of individual nAChR subunits (OK6-Gal4;UAS-dicer/UAS-RNAi) compared to UAS-RNAi/+ controls. Knockdown of α1–α6 and β2–β3 significantly reduces crawling path length. (C) Knockdown of individual nAChR subunits in a smaller subset of MNs (CQ-Gal4;27E09-Gal4) phenocopies the defects observed in (B), confirming consistency across independent drivers. (D) Representative crawling trajectories. Control larvae show normal movement; α7 knockdown larvae show no impairment, α1 knockdown larvae show mild defects, and β2 knockdown larvae show severe deficits. Violin plots show individual larvae (dots), with medians and IQRs indicated. Statistical significance was assessed using the Kruskal-Wallis test followed by Dunn’s post-hoc test (*P < 0.05, **P < 0.01, ****P < 0.0001). N > 39 larvae per group. Genotypes are provided in the SI Appendix.

Next, we used a more restricted MN driver, CQ-Gal4;27E09-Gal4, which targets a subset of MNs (five Type Ib and two Type Is MNs per hemisegment). This driver recapitulated the results from OK6-Gal4 knockdowns, reinforcing the importance of α1–α6, β2, and β3 subunits in supporting locomotion, while again showing no effect for α7 and β1 (**Fig. 2C**). These results underscore the essential role of specific nAChR subunits in mediating ionotropic excitation required for proper motor output.

To further probe the behavioral consequences of subunit knockdown on MN function, we analyzed two key parameters of peristaltic crawling: peristalsis efficiency (length traveled per peristalsis wave) and peristalsis duration (time per peristalsis wave). Knockdown of α1, α2, α4, α5, α6, β2, and β3 significantly reduced peristalsis efficiency, while α3, α7, and β1 knockdowns had no effect (**Fig. 3C** and **Movie S2**). Furthermore, peristalsis duration was significantly increased upon knockdown of all subunits except α7 and β1 (**Fig. 3D** and **Movie S2**). These results suggest that reduced nAChR expression impairs MN excitability, resulting in delayed activation or weakened output, which in turn slows the propagation or strength of peristaltic contractions.

**Figure 3.**
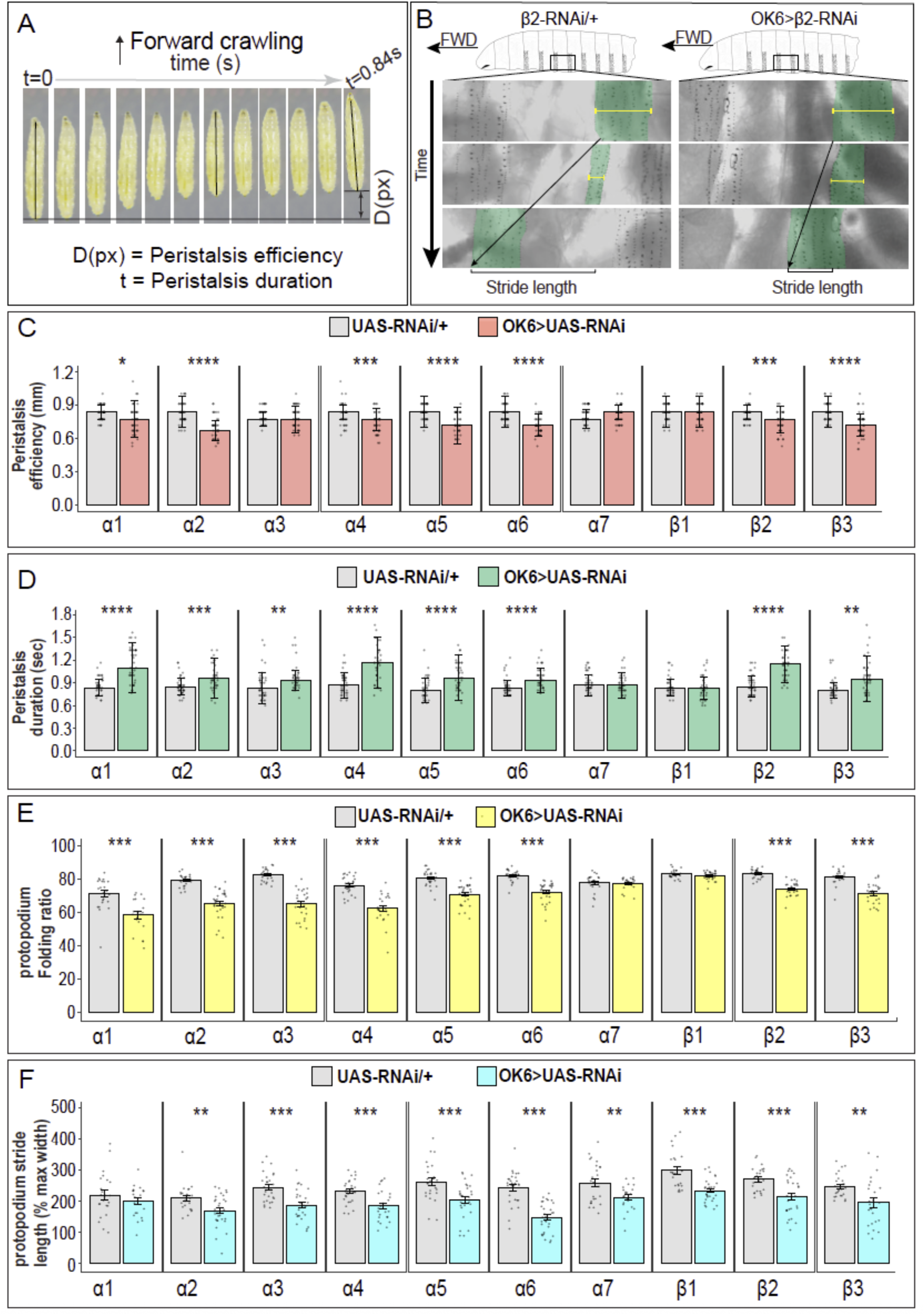
Single nAChR Subunit Knockdown in Motor Neurons Affects Crawling Parameters. **(A)** Representative time-lapse images of a single forward peristalsis in larval Drosophila. Peristalsis efficiency is defined as the distance traveled (D) per peristaltic wave, and peristalsis duration as the time (t) to complete one wave. **(B)** Stride length and protopodium folding visualized in control and β2 knockdown larvae. Knockdown of β2 severely reduces stride length and protopodium folding (yellow line). **(C)** Knockdown of α1, α2, α4, α5, α6, β2, and β3 significantly reduces peristalsis efficiency, while α3, α7, and β1 knockdowns do not. (D) Peristalsis duration is significantly increased with knockdown of α1, α2, α3, α4, α5, α6, β2, and β3. (E) Protopodium folding ratio is significantly reduced with knockdown of α1, α2, α3, α4, α5, α6, β2, and β3, but unaffected by α7 or β1 knockdowns. (F) Stride length is significantly reduced following knockdown of all subunits except α1. In (C–D), violin plots represent individual larvae; in (E–F), bar graphs show mean ± SEM with 2–3 protopodia per larva. Statistical significance was assessed using the Kruskal-Wallis test (C–D) and Student’s t-test (E–F). N > 39 larvae (C–D), N > 14 larvae (E–F). *P < 0.05, **P < 0.01, ****P < 0.0001. Genotypes are provided in the SI Appendix.

During crawling, larvae interact with the substrate using segmentally repeating structures that resemble primitive feet, termed protopodia (66-68). Each protopodium is composed of 6–8 rows of actin-rich denticles arranged across the ventroanterior part of each body segment. When crawling forward, the protopodium of the contracting segment enters a swing phase, folding and lifting off the ground, followed by propelling the segment forward, whereas the protopodia of non-contracting segments remain in the stance phase.

We thus investigated the influence of nAChR subunits on protopodium dynamics during crawling. We first quantified the protopodium folding ratio (contracted/resting width) and found that knockdown of α1–α6, β2, and β3 significantly reduced folding, while α7 and β1 knockdowns had no effect (**Fig. 3E** and **Movie S3**). We then assessed normalized stride length (displacement of the protopodium per peristaltic contraction wave) and found significant reductions upon knockdown of all subunits except α1 (**Fig. 3F** and **Movie S3**), suggesting broad involvement of nAChRs in this aspect of motor control.

Together, these functional assays reveal that nAChRs are critical regulators of larval locomotion. Subunits α1–α6, β2, and β3 contribute to multiple aspects of MN function, including peristalsis efficiency, timing, and protopodia dynamics. In contrast, α7 and β1 appear to play limited or dispensable roles in this context. These findings demonstrate a complex and finely tuned interplay among nAChR subtypes in shaping motor output and highlight the importance of cholinergic ionotropic signaling in driving coordinated locomotion in Drosophila larvae.

### Co-localization and Double Knockdown Analyses Identify nAChR Subunits That Localize to the Same Versus Distinct Postsynaptic Sites on Motor Neurons

Next, we sought to investigate the co-localization patterns of different nAChR subunit pairs at the synapse level in MN dendrites. For this, we utilized endogenously tagged subunits with distinct epitopes (**Fig. 4A**). Imaging was focused on the dorsal neuropil, where motor circuits are localized. To ensure the reliability of our co-localization analysis, we established thresholds using both positive and negative controls. Co-localization of the positive control, α6::HA and α6::OLLAS, showed over 98% overlap in the quantified puncta, validating the sensitivity of our parameters (**Fig. 4B**). For the negative control, we analyzed the overlap between Rdl::HA and α5::OLLAS. Rdl is a GABA-gated chloride channel, and we expected minimal overlap with the nAChR subunit α5 due to their distinct neurotransmitter systems. As anticipated, co-localization between Rdl::HA and α5::OLLAS was less than 2%, confirming the specificity of our co-localization criteria (**Fig. 4C**). These controls demonstrate the robustness of our methodology for assessing co-localization between nAChR subunits.

**Figure 4.**
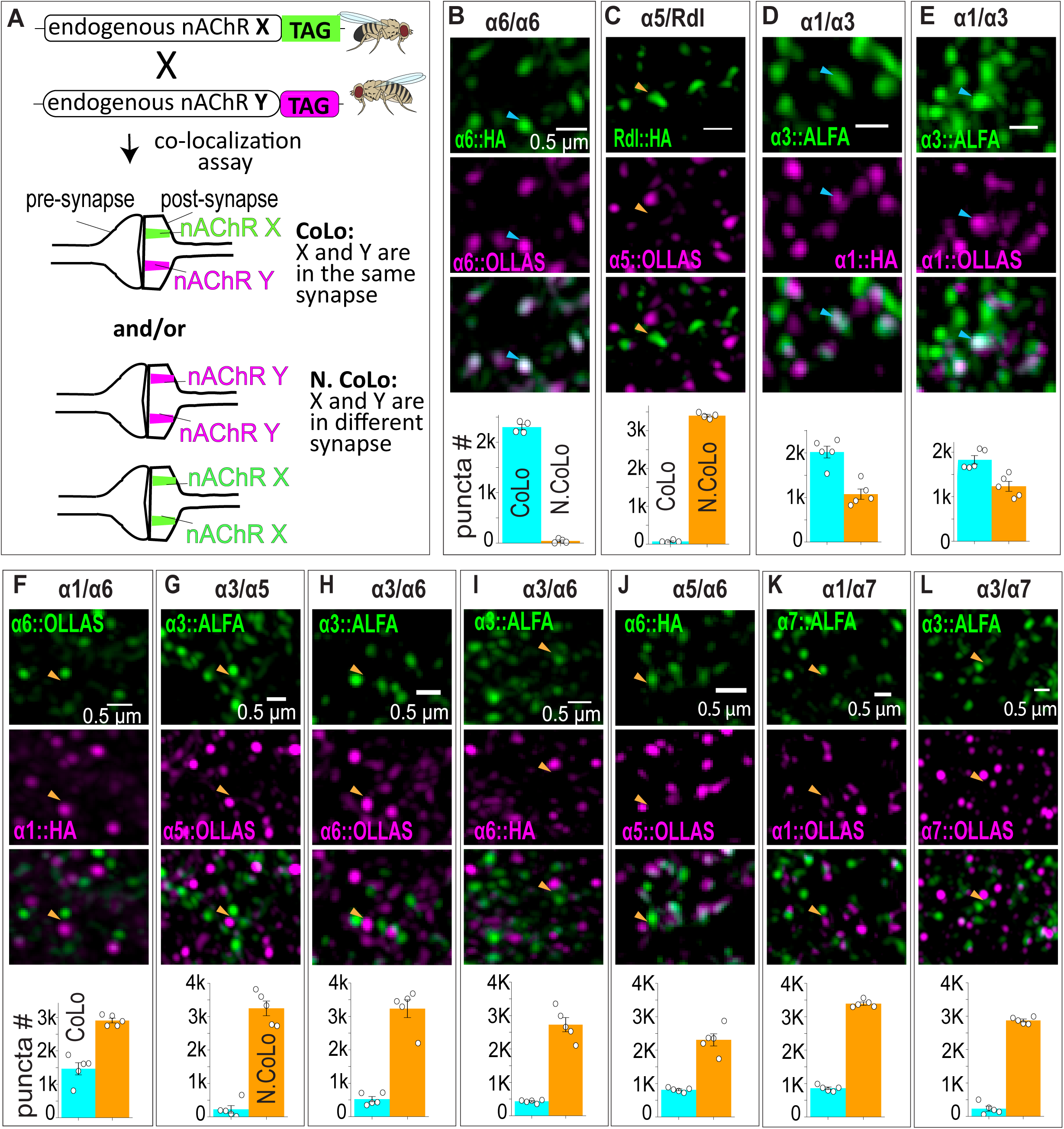
Most nAChR Subunits Are Spatially Segregated at Motor Neuron Synapses. (A) Schematic illustrating the strategy for assessing spatial relationships between endogenously tagged nAChR subunits. (B–C) Control experiments. (B) Positive control shows high colocalization between α6::HA and α6::OLLAS. (C) Negative control shows minimal colocalization between Rdl::HA and α5::OLLAS. (D–L) Colocalization patterns of nAChR subunit pairs. (D–E) High colocalization between α3::ALFA and α1::HA (D) and α1::OLLAS and α3::ALFA (E). (F–L) Predominantly non-colocalized patterns for α1::HA with α6::OLLAS (F), α3::ALFA with α5::OLLAS (G), α3::ALFA with α6::OLLAS (H), α3::ALFA with α6::HA (I), α5::OLLAS with α6::HA (J), α1::OLLAS with α7::ALFA (K), and α3::ALFA with α7::OLLAS (L). Cyan arrowheads indicate colocalized puncta; orange arrowheads indicate non-colocalized puncta. Bar graphs show individual brain measurements (n = 4–5), mean ± SEM. Genotypes are provided in the SI Appendix.

Building on these controls, we performed co-localization analyses for specific nAChR subunit pairings to determine their spatial organization within MN synapses. Double immunostaining with α3::ALFA and α1::HA revealed that α1 overlaps with α3 in over 65% of quantified puncta, indicating that α1 and α3 subunits may co-exist at the majority of synapses (**Fig. 4D**). Similar results were obtained using α3::ALFA and α1::OLLAS (**Fig. 4E**).

In contrast, co-localization assays for other subunit pairings exhibited substantially lower overlaps. For instance, double immunostaining of α1::HA with α6::OLLAS revealed only 36% co-localization, indicating that α1 and α6 are predominantly localized to different synapses (**Fig. 4F**). Similarly, α3::ALFA and α5::OLLAS showed minimal overlap, with only 7% of quantified puncta co-localized (**Fig. 4G**). Analyses involving α3::ALFA and α6::OLLAS revealed 14% co-localization (**Fig. 4H**), corroborated by a parallel assay using α3::ALFA and α6::HA (**Fig. 4I**). Co-localization between α5::OLLAS and α6::HA was observed in 26% of puncta (**Fig. 4J**), and α1::OLLAS with α7::ALFA exhibited 26% overlap (**Fig. 4K**). The pairing of α3::ALFA with α7::OLLAS showed minimal co-localization at 7% (**Fig. 4L**).

We next reasoned that for subunits belonging to distinct synapses, double knockdown in MNs should impair a broader range of cholinergic synapses and thus cause more severe locomotor defects than single knockdowns. In contrast, for subunits that predominantly co-localize at the same synapse, double knockdown would be less likely to exacerbate locomotor defects relative to single knockdowns. Our larval crawling assay results strongly support these hypotheses.

As expected, combined knockdown of the spatially segregated subunits α1/α6, α3/α5, α3/α6, and α5/α6 resulted in significantly more severe locomotor defects compared to individual knockdowns (**Fig. 5A–D**). Conversely, double knockdown of the co-localized pair α1/α3 did not lead to a more severe phenotype compared to the respective single knockdowns (**Fig. 5E**). Similarly, double knockdown of α1/α7 and α3/α7 also did not intensify locomotor defects relative to single knockdowns (**Fig. 5F–G**).

**Figure 5.**
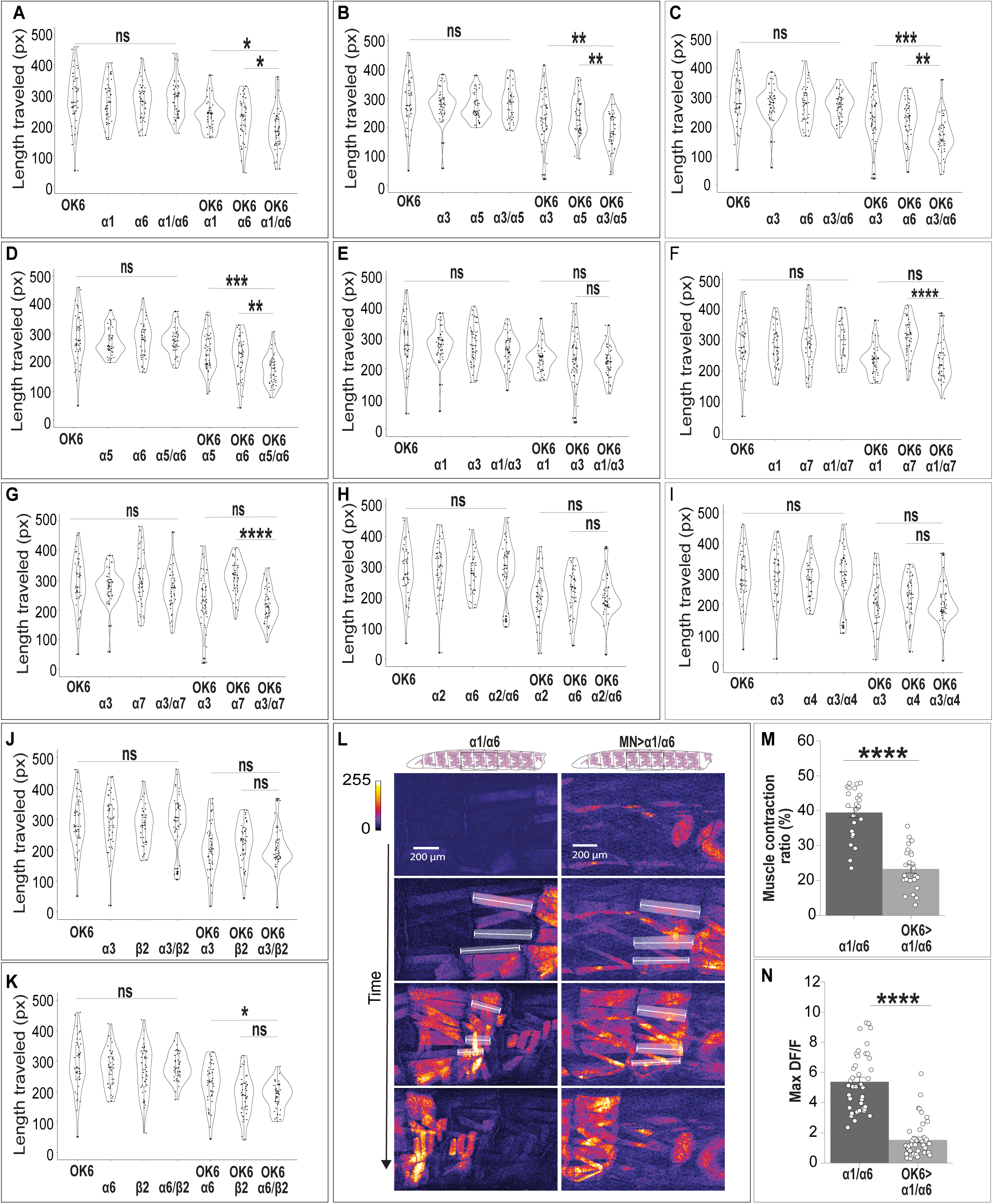
Simultaneous Knockdowns of Spatially Segregated nAChR Subunits Worsen Locomotion Beyond Single Subunit Loss. (A–K) Quantification of path length following single or dual nAChR subunit knockdowns in motor neurons. Dual knockdowns of spatially segregated subunits (α1/α6, α3/α5, α3/α6, α5/α6) cause significantly greater locomotor impairment compared to single knockdowns (A–D). Dual knockdown of spatially co-localized subunits (α1/α3) does not worsen defects (E). Similarly, α1/α7 (F), α3/α7 (G), α2/α6 (H), α3/α4 (I), and α3/β2 (J) dual knockdowns do not cause additive effects; α6/β2 (K) shows a trend without reaching significance. (L) Still images of GCaMP6f fluorescence and muscle contractions in control and α1/α6 dual knockdown larvae. (M–N) Quantification of muscle contraction ratio (M) and maximum ΔF/F calcium signals (N), both significantly reduced following α1/α6 dual knockdown. Violin plots (A–K) show individual larvae, with medians and IQRs indicated. Statistical significance was assessed using the Kruskal-Wallis test (A–K) and Student’s t-test (M–N). N > 39 larvae (A–K); n = 4–5 muscles per group (M–N). *P < 0.05, **P < 0.01, ****P < 0.0001; ns = not significant. Genotypes are provided in the SI Appendix.

We also examined additional subunit pairs for which endogenous co-localization data were not available. Double knockdown of α2/α6, α3/α4, and α3/β2 did not result in significantly enhanced defects relative to individual knockdowns, suggesting that these subunits likely co-exist at the same synapses (**Fig. 5H–J**). In contrast, knockdown of the α6/β2 pair resulted in a semi-significant worsening of the phenotype, suggesting that α6 and β2 may partially segregate or have complementary roles within a subset of MN synapses (**Fig. 5K**).

Together, these findings highlight the complex organization and functional diversity of nAChR subunits at cholinergic MN synapses. The stronger phenotypic effects observed upon double knockdown of subunits localized to different synapses underscore the importance of spatial segregation and functional specialization in cholinergic signaling. Conversely, the lack of enhanced defects following double knockdown of highly co-localized subunits suggests redundancy or cooperative action within shared synapses. Overall, our results provide critical insights into the architecture of nAChR subunits and their contributions to motor circuit function, paving the way for future studies aimed at dissecting their precise roles in synaptic physiology.

### MN-Specific Loss of α1 and α6 nAChR Subunits Impairs Muscle Activation and Contractile Force

To directly assess the impact of α1 and α6 subunit loss on motor output, we performed high-resolution muscle calcium imaging in intact larvae during forward crawling. In healthy larvae, peristaltic muscle contractions propagate from posterior to anterior segments, with muscles of a single segment at a time engaging to produce maximal contractile force, reflecting precise intersegmental coordination. In contrast, in **OK6 > α1/α6** larvae, muscle contractions were significantly less robust compared to controls (**Fig. 5L–N**). While muscles in control larvae demonstrated an average contraction amplitude of approximately 39%, those in **OK6 > α1/α6** larvae showed a marked decrease, averaging only 23% (**Fig. 5M**). This reduction in muscle contraction likely reflects impaired excitatory signaling due to knockdown of the α1 and α6 nAChR subunits. The inability to sustain normal muscle contractions likely contributes to the diminished peristalsis and locomotor deficits observed in these animals.

Furthermore, calcium responses were significantly reduced in α1/α6 knockdown larvae compared to controls. Control larvae exhibited significantly higher maximum ΔF/F values relative to **MN > α1/α6** larvae, suggesting that α1/α6 knockdown in MNs diminishes the maximum fluorescence change, consistent with lower muscle activation or excitability (**Fig. 5N**). These findings demonstrate that α1 and α6 nAChR subunits are critical for sustaining robust muscle activation during peristaltic locomotion. Their loss in MNs leads to impaired excitatory signaling, resulting in reduced contractile force and weakened motor output.

## Discussion

### Spatial and Functional Diversity of nAChR Subunits in Motor Circuits

In this study, we investigated the expression, function, and synaptic organization of nicotinic acetylcholine receptor (nAChR) subunits in *Drosophila* larvae, with a particular focus on motor neurons (MNs). We show that larval MNs express a broad array of subunits, including α1–α3, α5–α7, β1, and β2. Given that MNs must integrate temporally precise excitatory and inhibitory inputs to drive coordinated locomotion, this subunit diversity and synaptic specificity likely confer the precision required for motor output.

Our single RNAi knockdown experiments revealed that nAChR-mediated excitation is essential for larval crawling. Knockdown of most subunits—except α7 and β1—caused pronounced defects in peristaltic wave propagation, stride length, and protopodium dynamics. These findings suggest that MNs depend on a functionally diverse set of nAChRs to maintain robust motor performance.

To determine whether different nAChR subunits are assembled into the same or distinct postsynaptic domains, we combined co-localization analyses of endogenously tagged subunits with dual RNAi knockdowns. Subunits that showed low co-localization (e.g., α1/α6, α3/α6) produced additive or synergistic phenotypes when knocked down together, whereas strongly co-localized subunits (e.g., α1/α3) did not. These findings support a model in which MNs spatially segregate distinct nAChR pentamers to decode input from different cholinergic premotor neurons (PMNs). Such compartmentalization may enable input-specific responses depending on behavioral context—for example, distinguishing inputs from PMNs active during forward crawling (e.g., A27h), backward escape (e.g., A18b), or both (e.g., A18a).

### Receptor Compartmentalization May Be a Shared Principle Across Neural Circuits

This extensive receptor diversity and spatial organization is not unique to MNs but appears to be a conserved principle across *Drosophila* neural circuits. In the visual system, for example, T5 direction-selective neurons express multiple nAChR subunits (α1, α3, α4, α5, α7, β1) and the muscarinic receptor mAChR-B, while T4 neurons express α1, α5, and β1 (20-22). These receptors show highly stereotyped spatial distributions at synaptic sites. Just as MNs require precise input integration for coordinated locomotion, T4 and T5 neurons depend on specialized cholinergic signaling to compute motion direction. Notably, different nAChR subunits localize to distinct postsynaptic domains within T4 and T5 neurons. In T4 dendrites, nAChRα5 localizes to the middle domain to receive input from Mi1 and Tm3 neurons, while nAChRβ1 and α1 localize to distal dendrites where they form dendro-dendritic synapses with neighboring T4 neurons (20). In T5 dendrites, nAChRα5 predominantly localizes to synapses receiving input from Tm1, Tm2, and Tm4, whereas nAChRα7 associates primarily with Tm9 synapses. This molecular specificity mirrors the functional specialization of the presynaptic partners: Tm1, Tm2, and Tm4 exhibit transient responses to visual stimuli, while Tm9 provides sustained input (22). These observations support our broader model that neurons spatially segregate nAChR subtypes to decode input identity and temporal dynamics—whether in motor circuits or in the visual system.

### From Transcriptomics to Functional Specificity

Transcriptomic studies in vertebrates (69-71), adult flies (12), and larvae (62) support the notion that individual neurons co-express multiple nAChR subunits. However, transcriptomic data alone do not resolve where subunits localize at synapses. For example, larval MNs and adult male ejaculatory neurons express α1, α2, α3, β1, and β2, but our imaging data show that these subunits do not always co-localize. This highlights the need to validate transcriptomic predictions at the protein level and with subcellular resolution.

Furthermore, transcriptomic evidence for subunit compensation—for instance, upregulation of α5–α7 following α2 knockdown (12)—suggests that receptor composition can be dynamically remodeled. While this plasticity may complicate RNAi interpretation, it also underscores the adaptive flexibility of nAChR systems across developmental stages and behavioral demands.

Recent dual knockout studies support the notion that nAChR subunits exhibit both redundancy and specialization. For example, while α1 and β2 single mutants show mild or no wing inflation defects, their combined loss causes a severe phenotype (33). However, this same combination leads to greater reductions in climbing and lifespan than either mutant alone, supporting non-redundant behavioral functions (33). This finding aligns with our data showing that MN-specific subunit knockdowns produce distinct locomotor phenotypes, indicating minimal compensation during motor behavior.

### Context-Dependent Roles of nAChR Subunits

The organizational principles observed in *Drosophila* are echoed in vertebrate systems. Mammalian nAChRs, assembled from a larger set of subunits (α2–α10, β2–β4), form subtype-specific receptors like α4β2 and α7, each with distinct kinetic and pharmacological properties (72). These subtypes play essential roles in cognition, neurodevelopment, and attention, and exhibit dynamic spatial and temporal expression patterns (70, 72, 73).

Importantly, the requirement for specific subunits can vary with developmental stage. In mammals, a subunit may be critical in early development but dispensable in adults (9). Similarly, α7 is required for the adult escape reflex but not for larval crawling. In contrast, larval MN-specific knockdown of most subunits (except α7 and β1) impairs crawling, whereas adult null mutants for α1, α2, α3, and α6—but not α4, α5, α7, β2, or β3—display reduced climbing ability (10). These findings underscore the importance of temporospatial context when interpreting subunit function.

nAChRs also exhibit pleiotropic roles beyond the nervous system. In mammals, they are expressed in non-neuronal tissues where they regulate inflammation and tissue repair (74). In *Drosophila*, mutations in α1, β1, and β2 disrupt wing inflation and abdominal morphology (33, 75), suggesting potential non-neuronal roles in insects as well.

### Receptor-Aware Perspectives on Insecticide Sensitivity and Circuit Mapping

nAChRs are major targets of widely used insecticides such as neonicotinoids and spinosyns. These compounds overactivate cholinergic receptors, causing paralysis and death (76). Sensitivity to these toxins depends on subunit composition: α1 and β2 mutations confer resistance to neonicotinoids, while α6 is essential for spinosyn sensitivity (10, 77). Knockdown of α2 increases insecticide sensitivity in adults due to reduced expression of low-affinity subunits α5–α7 (12, 76). Our study provides a framework for predicting insecticide vulnerabilities based on MN-expressed subunits and their likely pairings. The observation that some subunits are co-expressed but not co-localized suggests that dual-targeting insecticides designed to engage distinct synapses may be more effective.

Although EM connectomes have greatly advanced our understanding of circuit architecture, they do not reveal the molecular components responsible for synaptic specificity (78, 79). Our data demonstrate that nAChR subunits are differentially localized even within single neurons, suggesting that distinct receptor assemblies decode input from specific presynaptic partners. Integrating molecular receptor mapping with anatomical connectomes will be essential to understanding circuit computation, behavioral function, and pharmacological targeting.

## Conclusion

Our findings reveal that *Drosophila* larval MNs employ a spatially and functionally diverse set of nAChR subunits to decode cholinergic inputs and drive coordinated locomotion. This work provides a framework for understanding how receptor diversity contributes to circuit logic and behavior.

Future efforts should explore how nAChRs interact with neuromodulators, glia, and structural scaffolds to regulate receptor localization and dynamics. High-resolution imaging, proteomic tagging, and integrated circuit mapping will be key to illuminating the principles of receptor-based computation in neural systems.

## Methods

Expression patterns of nAChR subunits were mapped in intact larvae using T2A-Gal4 lines and confirmed via immunohistochemistry with endogenously tagged subunits. MN-specific RNAi knockdown was achieved using two different MN driver lines, OK6-Gal4 and CQ-Gal4;27E09-Gal4, and the effects on larval crawling behavior were quantified using ImageJ and the wrMTrck plugin. Co-localization patterns of nAChRs in the neuropil, where MN dendrites reside, were analyzed using confocal imaging and object-based co-localization analysis in Imaris 10.0.1. Calcium imaging was conducted in intact larvae, with changes in GCaMP6s signals in muscles quantified using MATLAB code. Statistical analyses were performed using either Student’s t-test or the Kruskal-Wallis test followed by Dunn’s post-hoc test. Significance levels are indicated as ns (not significant), *P < 0.05, **P < 0.01, and ***P < 0.001. A detailed description of reagents and methods is provided in the SI Appendix.

## Author Contributions

Paste the author contributions here.

## Classification

BIOLOGICAL SCIENCES-Neuroscience

## Data Availability

All study data are included in the article and/or supporting information.

## Acknowledgments

We are grateful to the Bloomington Drosophila Stock Center, Shu Kondo, and S. Lawrence Zipursky for generously providing transgenic fly lines, and to Chihiro Hama for the anti-α6 antibody. We also thank Yuhan Huang and Lewis S. Sherer for their invaluable technical support, and Peter Newstein for providing the code used in the scRNA-seq data analysis. This research was funded by a grant from Texas A&M University.

