## Supporting information for "Motor Neurons Decode Cholinergic Inputs via Spatially Distinct nAChR Subunits to Drive Locomotion in Drosophila larvae"

**This PDF file includes:**

Supporting text (methods)
Figure S1
Table S1
Legends for Movie S1 to S3
SI References

**Other supporting materials for this manuscript include the following:**

Movies S1 to S3

### Supporting Information – methods

#### Fly culturing

Flies were maintained and reared on standard cornmeal, molasses and yeast based medium at 25 ± 1° C, ~50% humidity with a photoperiod of 12 hours light :12 hours night. For experiments involving RNAi, larvae were raised at 29 ± 1° C to increase RNAi efficiency. A complete list of fly stocks used in this study can be found in the key resource table (Tables **S1**).

#### Behavioral analyses and quantification

Wandering stage L3 larvae were selected from the cultured bottle and washed twice in distilled water. 6-10 larvae were then transferred to a 17 x 17 cm wide arena filled with 2% agarose using blunt forceps. Larval movement assay was adapted from a previously published paper (1). To ensure the arena surface was dry and smooth, a gentle brush was used to remove any remaining water droplets that could potentially disrupt larval locomotion. The arena was then placed under a digital camera, and following a 10 second acclimation period, the locomotor activity of the larvae was recorded for 1 minute at 30 frames per second using an iPhone mounted 30 cm from the agarose surface. The locomotor activity of the larvae was quantified using ImageJ software with the wrmTRck plugin with conditions optimized for the *Drosophila* larvae. First, a region of interest (ROI) was selected within the 60-second video to focus on the movement of the larvae. The wrmTRck plugin was then used to track the movement of larvae within the ROI, to measure the length traveled by each larva during the 1-minute recording period.

For the peristalsis efficiency and peristalsis duration assays, a two-layer 2% agarose was prepared in a 140 x 20 mm glass Petri dish. The base layer of agarose was poured into the glass and left to solidify. A groove mold was placed on top of the solidified base layer. An additional layer of agarose was then poured into the dish, covering the groove mold and allowed to solidify. The groove mold was carefully removed, leaving empty agarose grooves in the dish. The dish was placed on a sheet of paper with grids for reference. Larvae were placed into the empty agarose grooves using blunt forceps and a paintbrush. Videos were recorded using an iPhone mounted onto a ZEISS Stemi 305 microscope with similar setup as described previously (2). To evaluate peristalsis efficiency, the number of peristaltic waves required for the larvae to travel a length of 1 cm was manually counted. The length traveled per peristalsis was then determined by dividing the total length (10 mm) by the number of peristaltic waves. Peristalsis duration was assessed by manually counting the number of frames required for a peristaltic wave to propagate from the posterior body region to the anterior mouth hooks. The duration was then calculated by dividing the number of frames by the camera's frame rate to obtain time measurements.

**Protopodium swing-stance assay and quantification**

For the assessment of protopodia dynamics, third instar larvae were gently immobilized ventral side up on a 2% agarose gel pad to ensure consistency of orientation. A cover slip was carefully placed over the larva to maintain positioning without exerting excessive pressure. This preparation was then positioned on the stage of a confocal microscope equipped with brightfield imaging capabilities. Multiple peristalses were recorded as time series for each larva. Quantification of protopodium folding and displacement was done with custom MATLAB scripts. A linear ROI was placed on each protopodium, oriented perpendicular to the denticle bands and spanning across them. The length and position of the perpendicular line were adjusted frame by frame to account for protopodium movement and changes in width during a forward crawl. The protopodium folding ratio was determined by calculating the proportion of the protopodium width that was reduced during folding. This was done by subtracting the minimum width from the maximum width to obtain the width reduction, then dividing this value by the maximum width to normalize the result. To express the folding ratio as a percentage, the result was multiplied by 100. Protopodium displacement was calculated as the distance traveled by the protopodium ROI midpoint during each forward crawl normalized to the maximum protopodium width (in μm).

**Immunohistochemistry**

The late 3rd instar wandering larval brains were dissected in HL3.1 (hemolymph-like solution), fixed in 4% PFA (paraformaldehyde in 1x PBST) in a 24-well plate for 12 minutes at room temperature, and washed three times with PBST (0.3% Triton X-100 in 1x PBS) for 15 minutes each. Samples were then blocked overnight at 2 °C in PBST supplemented with 2% BSA (Fisher, BP1600-100), 1% normal donkey serum and 1% normal goat serum (Jackson ImmunoResearch Laboratories, 017-000-121 and 005-000-121). After blocking, brains were incubated in primary antibodies overnight at 2 °C. The primary antibodies were removed and washed three times in PBST for 15 minutes each. Brains were then incubated with secondary antibodies overnight at 2 °C. The secondary antibodies were removed, and the brains were washed three times in 0.3% PBST for 15 minutes each. Once washed, brains were mounted onto microscopic slides in Fluoromount-G (SouthernBiotech #0100-01) and stored at -20 °C for imaging purposes.

Primary antibodies : Rabbit anti-V5 (1:500, Cell Signaling Technology), Mouse anti-Brp/Nc82 (1:100, Developmental Studies Hybridoma Bank), Chicken anti-GFP (1:500, Invitrogen), Mouse anti-GFP (1:100, Developmental Studies Hybridoma Bank); Rabbit anti-GFP (1:500, Thermo Fisher Scientific), Mouse anti-HA (1:1000, BioLegend), Rat anti-HA (1:200, SigmaAldrich), Rat anti-OLLAS (1:100, Novus Biologicals), Rabbit anti-Dα6 (1:1000; from Mr.HAMA (3)), Atto 488 Alpaca anti-ALFA (1:500; Nanotag).

All secondary antibodies were purchased from Thermo Fisher Scientific and used at a working concentration of 1:200. The following antibodies were used: Alexa Fluor 405 Goat anti-Mouse, Alexa Fluor 488 Goat anti-Mouse, Alexa Fluor 488 Goat anti-Rabbit, Alexa Fluor 488 Donkey anti-Chicken, Alexa Fluor 555 Goat anti-Mouse, Alexa Fluor 555 Goat anti-Rat, Alexa Fluor 555 Goat anti-Rabbit, Alexa Fluor 594 Goat anti-Rat, Alexa Fluor 594 Goat anti-Rabbit, Alexa Fluor 647 Goat anti-Rat, Alexa Fluor 647 Donkey anti-Mouse, Alexa Fluor 647 Donkey anti-Rabbit.

**Image acquisition and quantification for co-localization**

Airyscan images of the dissected ventral nerve cord (VNC) in third instar wandering larval brains were acquired using a Zeiss LSM900 confocal microscope equipped with a 63× oil immersion objective (NA 1.4). Imaging was conducted in super-resolution (SR) mode with Zen Blue software. Raw image stacks were processed in Zen Blue software to construct Airyscan images using 3D Airyscan processing with automatic settings. To determine whether two subunits are co-localized within the same synapse, object-based co-localization analysis was performed on the dorsal neuropil region of the VNC, where synapses forming the motor circuit are located, using Imaris Bitplane 10.1. Separate surfaces were generated for each fluorescently labeled channel corresponding to the tagged subunits. The "Overlapped Volume to Surfaces" filter in Imaris was employed to calculate the degree of spatial overlap between the two subunit surfaces by applying threshold based on control samples. For statistical analysis, the number of co-localized and non-co-localized puncta for each subunit pair was quantified and represented graphically.

**Live imaging and quantification**

For live imaging, second- and third-instar larvae were washed with distilled water and placed on a 2% agarose pad positioned on a glass slide. A 2 mm x 40 mm coverslip was used to gently press the larvae into the agarose pad to immobilize them. A z-stack of the body wall was acquired using a 10x objective. For live calcium imaging of intact muscles in third-instar larvae, multiple peristaltic cycles were recorded as a time series using a similar setup, ensuring at least 40% of the muscle area was captured during imaging.

Muscle contraction was analyzed using a custom MATLAB script. A linear ROI was placed on each muscle, oriented parallel to the muscle of interest. The position and length of the ROI were adjusted frame by frame to account for changes in muscle width during forward crawling. Contraction was quantified as the proportion of muscle width reduced during the crawl, calculated by subtracting the minimum width from the maximum width to determine the reduction, then normalizing this value by dividing by the maximum width. The contraction ratio was expressed as a percentage by multiplying the result by 100.

**Single-cell RNA-seq (scRNA-seq) data analysis**

Publicly available scRNA-seq data were obtained from the NCBI Gene Expression Omnibus (GEO) under accession number GSE235231 (4). The raw count matrix and accompanying metadata were processed using the Seurat package (v4.3.0) in R. Transcriptional clusters were annotated with motor nerve bundle identities based on anatomical information provided in the original publication. Clusters corresponding to the same motor bundle were grouped under a unified label for visualization. Expression of nicotinic acetylcholine receptor (nAChR) subunits was analyzed across these anatomically defined groups. A DotPlot was generated to display average log-normalized expression (dot color) and the proportion of expressing cells (dot size).

**Figure preparation**

Images in figures were prepared as 3D projections in FIJI (ImageJ 1.54g) and assembled using Adobe Illustrator or Adobe Photoshop. Schematics were drawn in BioRender.

**Statistical analysis**

Statistics were performed using a combination of Microsoft Excel, MATLAB (MathWorks), R-Studio, and Python (Jupyter software). For data involving larval movement assay, peristalsis efficiency, and peristalsis duration assay Kruskal-Wallis Test with Dunn's multiple comparison was done. For data involving protopodium swing-stance assay, and muscle contraction Student's t test was done. Levels of significance were established at *: $p<.05$, **: $p<0.01$, ***: $p<0.001$, ****$P < 0.0001$. All figures depict data in bar plots or violin plots with individual values. All other pertinent information, including sample size, statistical test used, and variance can be found in the figure legends or labelled within the figure.

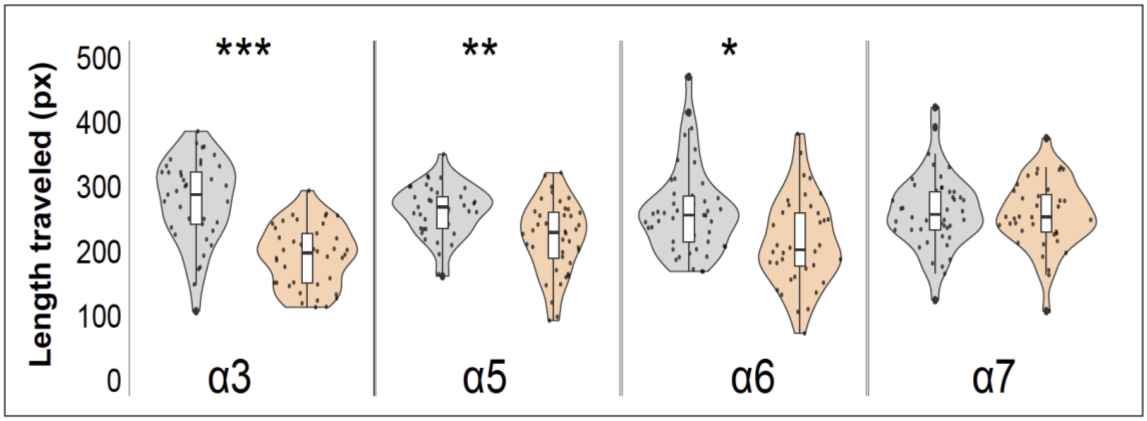


**Supplementary Figure 1. MN-specific knockdown of the same nAChR subunits using independent RNAi constructs produces consistent behavioral outcomes, ruling out RNAi off-target effects.** Quantification of total crawling path length in larvae with motor neuron (MN)-specific knockdown of nAChR subunits using OK6-Gal4;UAS-dicer/UAS-RNAi compared to UAS-RNAi/+ controls. Knockdown of α3, α5, and α6 significantly reduces locomotion, while knockdown of α7 has no observable effect on crawling behavior. These results are consistent with the findings presented in **Figure 2** of the main text, where different RNAi lines were used to selectively knock down the same subunits in MNs. Individual data points represent the crawling path length of a single larva. Data are presented as violin plots showing the distribution of path lengths, with the median and interquartile range (IQR) indicated within each plot to illustrate data variability. Statistical significance was assessed using the Kruskal-Wallis test followed by Dunn's post-hoc test (*$P < 0.05$, **$P < 0.01$, ****$P < 0.0001$). Sample sizes are $N > 39$ larvae per group.

**Tables S1**: Key Resources Table

| **Reagent type (species) or resource** | **Designation** | **Source or reference** | **Additional information** |
|---|---|---|---|
| Genetic Reagent (*Drosophila melanogaster*) | *UAS-myr-GFP; 44H10-LexA-OP-mCherry* | This study | Used in **Figure 1**. This was crossed to *T2A-Gal4* lines and MNs were imaged in intact offspring larvae. |
| Genetic reagent (*Drosophila melanogaster*) | *(Mi(4)nAChRalpha1[MI00453-TG4.0]/TM3, Sb[1] Ser[1])* | BDSC #66780 | Used in **Figure 1**. This was crossed to *10×UAS-myr::GFP* and MNs were imaged in intact offspring larvae. |
| Genetic reagent (*Drosophila melanogaster*) | *w[*]; TI(4)CG6844[PGxRF3-1]* | (5) | Used in **Figure 1**. This was crossed to *10×UAS-myr::GFP* and MNs were imaged in intact offspring larvae. |
| Genetic reagent (*Drosophila melanogaster*) | *w[*]; TI(6)CG2302[PGxRF3-1]* | (5) | Used in **Figure 1**. This was crossed to *10×UAS-myr::GFP* and MNs were imaged in intact offspring larvae. |
| Genetic reagent (*Drosophila melanogaster*) | *y[1] w[*]; Mi(6)nAChRalpha5[MI13859-TG4.2]/SM6a* | BDSC #76755 | Used in **Figure 1**. This was crossed to *10×UAS-myr::GFP* and MNs were imaged in intact offspring larvae. |
| Genetic reagent (*Drosophila melanogaster*) | *y[1] w[*]; Mi{Trojan-GAL4.1}(6)nAChRalpha6[MI01466-TG4.1]* | BDSC #76137 | Used in **Figure 1**. This was crossed to *10×UAS-myr::GFP* and MNs were imaged in intact offspring larvae. |
| Genetic reagent (*Drosophila melanogaster*) | *y[1] w[*] Mi{Trojan-GAL4.1}nAChRalpha7[MI12545-TG4.1]* | BDSC #77828 | Used in **Figure 1**. This was crossed to *10×UAS-myr::GFP* and MNs were imaged in intact offspring larvae. |
| Genetic reagent (*Drosophila Melanogaster*) | *w[*]; TI{T2A-GAL4}CG11348[RA-PGxRF3-1]* | (5) | Used in **Figure 1**. This was crossed to *10×UAS-myr::GFP* and MNs were imaged in intact offspring larvae. |
| Genetic reagent (*Drosophila Melanogaster*) | *w[*]; TI{T2A-GAL4}CG6798[RA-PGxRF3-2]/TM3* | (5) | Used in **Figure 1**. This was crossed to *10×UAS-myr::GFP* and MNs were imaged in intact offspring larvae. |
| Genetic reagent (*Drosophila Melanogaster*) | *w[*]; TI{T2A-GAL4}CG11822[PGxRF3-1]/CyO* | (5) | Used in **Figure 1**. This was crossed to *10×UAS-myr::GFP* and MNs were imaged in intact offspring larvae. |

| | | | |
|---|---|---|---|
| Genetic reagent (*Drosophila Melanogaster)* | *RRa-Gal4, UAS-myr::GFP* | This study | Used in **Figure 1**. This line was crossed to endogenously tagged nAChR lines, and the VNC of offspring larvae was imaged. |
| Genetic reagent (*Drosophila Melanogaster)* | *Da1-GFP11-3xHA* | (7) | Used in **Figure 1**. This line was crossed to *RRa-Gal4, UAS-myr::GFP* and, the VNC of offspring larvae was imaged. |
| Genetic reagent (*Drosophila Melanogaster)* | *Da6-GFP11-3xHA* | (7) | Used in **Figure 1**. This line was crossed to *RRa-Gal4, UAS-myr::GFP* and, the VNC of offspring larvae was imaged. |
| Genetic reagent (*Drosophila Melanogaster)* | *w;; nAChRβ1-KDRT-smGdP-10xHA* | (6) | Used in **Figure 1**. This line was crossed to *RRa-Gal4, UAS-myr::GFP* and, the VNC of offspring larvae was imaged. |
| Genetic reagent (*Drosophila Melanogaster)* | *94G06-Gal4; UAS-FLP* | This study | Used in **Figure 1**. This line was crossed to conditionally tagged nAChR lines, and the VNC of offspring larvae was imaged. |
| Genetic reagent (*Drosophila Melanogaster)* | *UAS-FRT-STOP-FRT-myrGFP-2A-KDRPEST; nAChRα1-KDRT-STOP-KDRT-smGdP-10xOllas* | (6) | Used in **Figure 1**. This line was crossed to *94G06-Gal4; UAS-FLP*, and the VNC of offspring larvae was imaged. |
| Genetic reagent (*Drosophila Melanogaster)* | *UAS-FRT-STOP-FRT-myrGFP-2A-KDRPEST; nAChRα5-KDRT-STOP-KDRT-smGdP-10xOllas* | (6) | Used in **Figure 1**. This line was crossed to *94G06-Gal4; UAS-FLP*, and the VNC of offspring larvae was imaged. |
| Genetic reagent (*Drosophila Melanogaster)* | *UAS-FRT-STOP-FRT-myrGFP-2A-KDRPEST; nAChRα6-KDRT-STOP-KDRT-smGdP-10xOllas, UAS-KDR* | (6) | Used in **Figure 1**. This line was crossed to *94G06-Gal4; UAS-FLP*, and the VNC of offspring larvae was imaged. |
| Genetic reagent (*Drosophila Melanogaster)* | *UAS-FRT-STOP-FRT-myrGFP-2A-KDRPEST; nAChRβ1-KDRT-STOP-KDRT-smGdP-10xHA/TM6B* | (6) | Used in **Figure 1**. This line was crossed to *94G06-Gal4; UAS-FLP*, and the VNC of offspring larvae was imaged. |
| Genetic reagent (*Drosophila melanogaster*) | $w^{1118}$ | BDSC | Crossed as control for nAChR knockdown experiments |

| | | | |
|---|---|---|---|
| Genetic reagent (*Drosophila melanogaster*) | *OK6-Gal4* | BDSC #64199 | MN specific nAChR knockdown. **Figure S1**. This line was crossed to *nAChR-RNAi* lines and locomotion was examined in offspring larvae. |
| Genetic reagent (*Drosophila melanogaster*) | *UAS-Dcr-2* | BDSC #24651 | This line was crossed with *OK6-Gal4*, then with VALIUM 10 *nAChR-RNAi* lines. |
| Genetic reagent (*Drosophila melanogaster*) | *OK6-Gal4 ; UAS-Dcr-2* | This study | Used in **Figures 2-3,5.**This line was crossed to VALIUM 10 *nAChR-RNAi* lines and locomotion was examined in offspring larvae. |
| Genetic reagent (*Drosophila melanogaster*) | *UAS-α1-RNAi* | BDSC #28688 | Used in **Figures 2-3**. This line was crossed to *OK6-Gal4 ; UAS-dicer* and locomotion was examined in offspring larvae. |
| Genetic reagent (*Drosophila melanogaster*) | *UAS-α2-RNAi* | BDSC #27493 | Used in **Figures 2-3**. This line was crossed to lines *OK6-Gal4 ; UAS-dicer* and locomotion was examined in offspring larvae. |
| Genetic reagent (*Drosophila melanogaster*) | *UAS-α3-RNAi* | BDSC #27671 | Used in **Figures 2-3**. This line was crossed to lines *OK6-Gal4 ; UAS-dicer* and locomotion was examined in offspring larvae. |
| Genetic reagent (*Drosophila melanogaster*) | *UAS-α3-RNAi* | BDSC #61225 | Used in **Figure S1**. This line was crossed to lines *OK6-Gal4,* and locomotion was examined in offspring larvae. |
| Genetic reagent (*Drosophila melanogaster*) | *UAS-Dα4-RNAi* | BDSC #31985 | Used in **Figures 2-3**. This line was crossed to lines *OK6-Gal4 ; UAS-dicer*, and locomotion was examined in offspring larvae. |
| Genetic reagent (*Drosophila melanogaster*) | *UAS-α5-RNAi* | BDSC #25943 | Used in **Figures 2-3**. This line was crossed to lines *OK6-Gal4 ; UAS-dicer,* and locomotion was examined in offspring larvae. |
| Genetic reagent (*Drosophila melanogaster*) | *UAS-α5-RNAi* | BDSC #77418 | Used in **Figure S1**. This line was crossed to lines *OK6-Gal4,* and locomotion was examined in offspring larvae. |
| Genetic reagent (*Drosophila melanogaster*) | *UAS-α6-RNAi* | BDSC #57818 | Used in **Figure S1**. This line was crossed to lines *OK6-Gal4*, and locomotion was examined in offspring larvae. |

| | | | |
|---|---|---|---|
| Genetic reagent (*Drosophila melanogaster*) | *UAS-a6-RNAi* | BDSC #52885 | Used in **Figures 2-3**. This line was crossed to lines *OK6-Gal4; UAS-dicer,* and locomotion was examined in offspring larvae. |
| Genetic reagent (*Drosophila melanogaster*) | *UAS-a7-RNAi* | BDSC #27251 | Used in **Figures 2-3**. This line was crossed to lines *OK6-Gal4,* and locomotion was examined in offspring larvae. |
| Genetic reagent (*Drosophila melanogaster*) | *UAS-α7-RNAi* | BDSC #51049 | Used in **Figure S1**. This line was crossed to lines *OK6-Gal4,* and locomotion was examined in offspring larvae. |
| Genetic reagent (*Drosophila melanogaster*) | *UAS-β1-RNAi* | BDSC #31883 | Used in **Figures 2-3**. This line was crossed to lines *OK6-Gal4; UAS-dicer,* and locomotion was examined in offspring larvae. |
| Genetic reagent (*Drosophila melanogaster*) | *UAS-β2-RNAi* | BDSC #28038 | Used in **Figures 2-3**. This line was crossed to lines *OK6-Gal4; UAS-dicer,* and locomotion was examined in offspring larvae. |
| Genetic reagent (*Drosophila melanogaster*) | *UAS-β3-RNAi* | BDSC #25927 | Used in **Figures 2-3**. This line was crossed to lines *OK6-Gal4; UAS-dicer,* and locomotion was examined in offspring larvae. |
| Genetic reagent (*Drosophila melanogaster*) | *α6-GFP11-3xHA nAChRα6-KDRT-smGdP-10xOLLAS* | This study | Used as a control in **Figure 4** to perform co-localization analysis. |
| Genetic reagent (*Drosophila melanogaster*) | *Rdl-KDRT-smGdP-10xHA nAChRα5-KDRT-smGdP-10xOLLAS* | This study | Used as a control in **Figure 4** to perform co-localization analysis. |
| Genetic reagent (*Drosophila melanogaster*) | *nAChRα3-KDRT-1xALFA α1-GFP11-3xHA* | This study | Used in **Figure 4** to perform co-localization analysis. |
| Genetic reagent (*Drosophila melanogaster*) | *nAChRα6-KDRT-smGdP-10xOLLAS α1-GFP11-3xHA* | This study | Used in **Figure 4** to perform co-localization analysis. |
| Genetic reagent (*Drosophila melanogaster*) | *nAChRα3-KDRT-1xALFA AChRα5-KDRT-smGdP-10xOLLAS* | This study | Used in **Figure 4** to perform co-localization analysis. |

| | | | |
|---|---|---|---|
| Genetic reagent (*Drosophila melanogaster*) | *nAChRα3-KDRT-1xALFA nAChRα6-KDRT-smGdP-10xOLLAS* | This study | Used in **Figure 4** to perform co-localization analysis. |
| Genetic reagent (*Drosophila melanogaster*) | *nAChRα5-KDRT-smGdP-10xOLLAS α6-GFP11-3xHA* | This study | Used in **Figure 4** to perform co-localization analysis. |
| Genetic reagent (*Drosophila melanogaster*) | *nAChRα3-KDRT-1xALFA nAChRα1-KDRT-smGdP-10xOLLAS* | This study | Used in **Figure 4** to perform co-localization analysis. |
| Genetic reagent (*Drosophila melanogaster*) | *nAChRα3-KDRT-1xALFA α6-GFP11-3xHA* | This study | Used in **Figure 4** to perform co-localization analysis. |
| Genetic reagent (*Drosophila melanogaster*) | *nAChRα7-KDRT-1xALFA α1-GFP11-3xHA* | This study | Used in **Figure 4** to perform co-localization analysis. |
| Genetic reagent (*Drosophila melanogaster*) | *nAChRα3-KDRT-1xALFA nAChRα7-KDRT-smGdP-10xOLLAS* | This study | Used in **Figure 4** to perform co-localization analysis. |
| Genetic reagent (*Drosophila melanogaster*) | *UAS-α1-RNAi ; UAS-α3-RNAi* | This study | Used in **Figure 5**. This line was crossed to lines *OK6-Gal4; UAS-dicer,* and locomotion was examined in offspring larvae. |
| Genetic reagent (*Drosophila melanogaster*) | *UAS-α1-RNAi ; UAS-α6-RNAi* | This study | Used in **Figure 5**. This line was crossed to lines *OK6-Gal4; UAS-dicer,* and locomotion was examined in offspring larvae. |
| Genetic reagent (*Drosophila melanogaster*) | *UAS-α3-RNAi ; UAS-α5-RNAi* | This study | Used in **Figure 5**. This line was crossed to lines *OK6-Gal4; UAS-dicer,* and locomotion was examined in offspring larvae. |
| Genetic reagent (*Drosophila melanogaster*) | *UAS-α3-RNAi ; UAS-α6-RNAi* | This study | Used in **Figure 5**. This line was crossed to lines *OK6-Gal4; UAS-dicer,* and locomotion was examined in offspring larvae. |
| Genetic reagent (*Drosophila melanogaster*) | *UAS-α5-RNAi ; UAS-α6-RNAi* | This study | Used in **Figure 5**. This line was crossed to lines *OK6-Gal4; UAS-dicer,* and locomotion was examined in offspring larvae. |

| | | | |
|---|---|---|---|
| Genetic reagent (*Drosophila melanogaster*) | *UAS-α1-RNAi ; UAS-α7-RNAi* | This study | Used in **Figure 5**. This line was crossed to lines *OK6-Gal4; UAS-dicer,* and locomotion was examined in offspring larvae. |
| Genetic reagent (*Drosophila melanogaster*) | *UAS-α3-RNAi ; UAS-α7-RNAi* | This study | Used in **Figure 5**. This line was crossed to lines *OK6-Gal4; UAS-dicer,* and locomotion was examined in offspring larvae. |
| Genetic reagent (*Drosophila melanogaster*) | *UAS-α3-RNAi ; UAS-α4-RNAi* | This study | Used in **Figure 5**. This line was crossed to lines *OK6-Gal4; UAS-dicer,* and locomotion was examined in offspring larvae. |
| Genetic reagent (*Drosophila melanogaster*) | *UAS-α3-RNAi ; UAS-β2-RNAi* | This study | Used in **Figure 5**. This line was crossed to lines *OK6-Gal4; UAS-dicer,* and locomotion was examined in offspring larvae. |
| Genetic reagent (*Drosophila melanogaster*) | *UAS-α2-RNAi ; UAS-α6-RNAi* | This study | Used in **Figure 5**. This line was crossed to lines *OK6-Gal4; UAS-dicer,* and locomotion was examined in offspring larvae. |
| Genetic reagent (*Drosophila melanogaster*) | *UAS-α3-RNAi ; UAS-β2-RNAi* | This study | Used in **Figure 5**. This line was crossed to lines *OK6-Gal4; UAS-dicer,* and locomotion was examined in offspring larvae. |
| Genetic reagent (*Drosophila melanogaster*) | *OK6-Gal4, UAS-Dcr-2; 44H10::GCaMP6f* | This study | Used in **Figure 5**. This line was crossed with *UAS-α6-RNAi; UAS-α1-RNAi*, and the muscles of the offspring larvae were imaged. |

**Movie S1.** nAChR knockdown in MNs affects larval locomotion**.** Video imaging of *Dα2-RNAi* alone and *OK6-Gal4 > Dα2-RNAi*. Dα2 knockdown in MNs significantly impairs length traveled by larvae.

**Movie S2.** nAChR knockdown in MNs affects peristalsis efficiency and duration. Video imaging of *Dβ2-RNAi* alone and *OK6-Gal4 > Dβ2-RNAi*. Dβ2 knockdown in MNs affects both peristalsis efficiency and duration.

**Movie S3.** nAChR knockdown in MNs affects stride length and protopodia folding ratio. Video imaging of *Dβ2-RNAi* alone and *OK6-Gal4 > Dβ2-RNAi*. Dβ2 knockdown in MNs affects both protopodia folding ratio and stride length.
